# pADP-ribosylation Regulates the Cytoplasmic Localization, Cleavage and Pro-apoptotic Function of HuR

**DOI:** 10.1101/2023.08.07.552262

**Authors:** Kholoud Ashour, Derek Hall, Souad Mubaid, Sandrine Busque, Xian Jin Lian, Jean-Philippe Gagné, Shahryar Khattak, Sergio Di Marco, Guy G. Poirier, Imed-Eddine Gallouzi

## Abstract

HuR (ElavL1) is one of the main posttranscriptional regulators that determines cell fate. While the role of HuR in apoptosis is well-established, the posttranslational modifications that governs this function remain elusive. In this study we show that PARP1/2-mediated poly(ADP)-ribosylation (PARylation) is instrumental in the pro-apoptotic function of HuR. During apoptosis or in cells depleted of PARP1/2 enzymes, a substantial reduction in HuR PARylation is observed. This results in the cytoplasmic accumulation and the cleavage of HuR, both of which are essential events for apoptosis. These effects are mediated by a pADP-ribose (PAR) binding motif within the HuR-HNS region (HuR PAR Binding Site (HuR-PBS)). Under normal conditions, the association of HuR-PBS with PAR is responsible for the nuclear retention of HuR. Mutations within this motif prevents the binding of HuR to its import factor TRN2, leading to its cytoplasmic accumulation and cleavage. Collectively, our findings underscore the role of PARylation in controlling the proapoptotic function of HuR, offering insight into the mechanism by which PARP1/2 enzymes regulate cell fate and adaptation to various assaults.

## Introduction

The RNA-binding protein HuR (Human antigen R) is a ubiquitously expressed protein. It contains three highly conserved RNA binding domains (RBD) also known as RNA recognition motifs (RRM; RRM1-3), and a hinge region between RRMs 2 and 3 that serves as the HuR Nucleocytoplasmic Shuttling (HNS) domain. HuR associates with many mRNA targets by binding mainly to the AU-rich elements (AREs) located in their 3′-untranslated region (UTR) thereby regulating their stability, translation, and/or subcellular localization (Grammatikakis *et al*, 2017b; Srikantan & Gorospe, 2012). HuR is a multifunctional protein that is known to be involved in many cellular processes, including cell proliferation and differentiation as well as apoptosis. Our laboratory, as well as others, have previously demonstrated that HuR is required for both pro-survival and pro-apoptotic processes (Lal *et al*, 2005; Mazroui *et al*, 2008; Von Roretz *et al*, 2013). It is known that in response to mild, sub-lethal stress, HuR modulates the expression of various pro-survival messages such as prothymosin α(Lal *et al*., 2005). However, when the stress is lethal, HuR shifts its function and modulates the expression of many pro-apoptotic factors such as caspase-9(Von Roretz *et al*., 2013). Our group has previously shown that this shift in function is promoted by the caspase 3/7-mediated cleavage of HuR (Von Roretz *et al*., 2013). Indeed, under apoptotic conditions, HuR translocates to the cytoplasm where it undergoes caspase-dependent cleavage at the aspartate (D)226 residue, thereby generating two cleavage products, HuR-CP1 (24 kDa) and HuR-CP2 (8 kDa). This event results in the cytoplasmic accumulation of HuR, where HuR mediates its pro-apoptotic function (Von Roretz *et al*., 2013).

While HuR is mainly localized to the nucleus under normal conditions, it has the ability to shuttle between the nucleus and the cytoplasm in response to various stimuli. This translocation is mediated via its interaction with protein partners, such as the nuclear export factor PHAPI (also known as pp32) and the nuclear import factor Transportin-2 (TRN2) (Mazroui *et al*., 2008; von Roretz *et al*, 2011b). It has been established that HuR and its cleavage products can be involved in apoptosis via their selective interaction with these protein partners. This interaction defines how HuR modulates the switch of two opposing processes: cell survival and cell death. PHAPI, a well-known HuR ligand, is an important activator of the apoptosome formation. In response to lethal stimuli, HuR/PHAPI complex is exported from the nucleus to the cytoplasm where both HuR and PHAPI exert their pro-apoptotic function(Mazroui *et al*., 2008; von Roretz *et al*., 2011b). The cytosolic cleavage of HuR facilitates the release of PHAPI mediating the activation of apoptosome-formation. Moreover, we also showed that HuR-CP2 selectively associates with PHAPI while HuR-CP1 interacts with TRN2, resulting in the decreased association of TRN2 with HuR. The binding of HuR-CP1 to TRN2, therefore, blocks the reimport of HuR into the nucleus leading to the accumulation of HuR in the cytoplasm, advancing apoptosis(Mazroui *et al*., 2008; von Roretz *et al*., 2011b).

The function of HuR, as well as other RBPs, has been shown to be primarily regulated through posttranslational modifications, such as phosphorylation, methylation and more recently poly (ADP-ribosyl) ation (PARylation)(Grammatikakis *et al*, 2017a; Ke *et al*, 2017). The process of PARylation, whereby poly-ADP-ribose (PAR) polymers are generated by PAR polymerase enzymes (PARPs), is known, in addition to its fundamental role in DNA repair, to be involved in apoptosis (Pleschke *et al*, 2000; Wei & Yu, 2016). The PARP family of proteins consist of seventeen members that are known to be involved in several cellular processes including PARP1, PARP2, PARP5a (TNKS1), and PARP5b (TNKS2)(Amé *et al*, 2004; Richard *et al*, 2022). The most characterised and well-studied enzyme of the PARP family is PARP1. The catalytic activity of PARP1 is normally initiated in response to a break in the DNA strand. When the DNA damage is mild and manageable, PARP1 detects and recruits DNA damage response factors to repair the damage and hereby acts as a cell survival factor. However, when the damage is irreparable, PARP1 is cleaved in the nucleus in a caspase-dependent manner by caspase 3 and 7, thereby leading to apoptosis (Diamantopoulos *et al*, 2014; Mashimo *et al*, 2021). Ke *et al*. have shown that the PARylation of HuR, via PARP1, regulates its localization and function during inflammation (Ke *et al*., 2017). They demonstrated that, in response to Lipopolysaccharide (LPS) exposure, PARP1 depletion/inhibition decreased the stability of mRNA from proinflammatory gene including *Cxcl2* (Ke *et al*., 2017). While PARPs can mediate the covalent PARylation of target proteins at specific residues, the catalyzed PAR chain can also bind in a non-covalent manner with proteins that contain a conserved PAR-binding motif (PBM) (Reber & Mangerich, 2021). Generally, this motif consists of loosely conserved sequence of hydrophobic and basic amino acids which are often found to overlap with important functional domains such as DNA and/or RNA binding domains, exerting regulatory function within the cell (Reber & Mangerich, 2021). Although PARylation of HuR is known to affect its function, the relevance of HuR binding to PAR on its function during apoptosis remains elusive.

Towards this end, in this study, we investigate the role of PARylation in the apoptotic function of HuR. We identified a PAR-binding motif (PBM) located in the HNS of HuR. Disruption of this motif results in the cytoplasmic localization of HuR and cell death. We demonstrate, thus, that the interaction of PAR with HuR, through the PBM, plays a key role in mediating the localization and function of HuR during apoptosis.

## Results

### PARP1 and PARP2 regulate the cytoplasmic localization, cleavage, and pro-apoptotic function of HuR

In order to establish the role of PARylation in the function of HuR during apoptosis we assessed, as a first step, whether HuR is associated with PAR polymers in HeLa cells treated, over a three-hour period, with staurosporine (STS), a well-known apoptotic inducer. We have previously demonstrated, as described in (Mazroui *et al*., 2008) and shown in (Fig. 1A) that the treatment of these cells with STS for 3h hours induces both the cleavage of HuR and PARP1. We show, under these conditions, by performing co-immunoprecipitation experiments, that although PAR associates to HuR in untreated cells, this interaction decreases when the cells were treated with STS for up to 3h (Fig.1B). The decreased interaction of HuR with PAR, interestingly, appears to coincide with the cleavage of PARP1 under these conditions (Fig. 1A) suggesting that PARP1 is involved in mediating the interaction of HuR with PAR in untreated cells.

**Figure 1:**
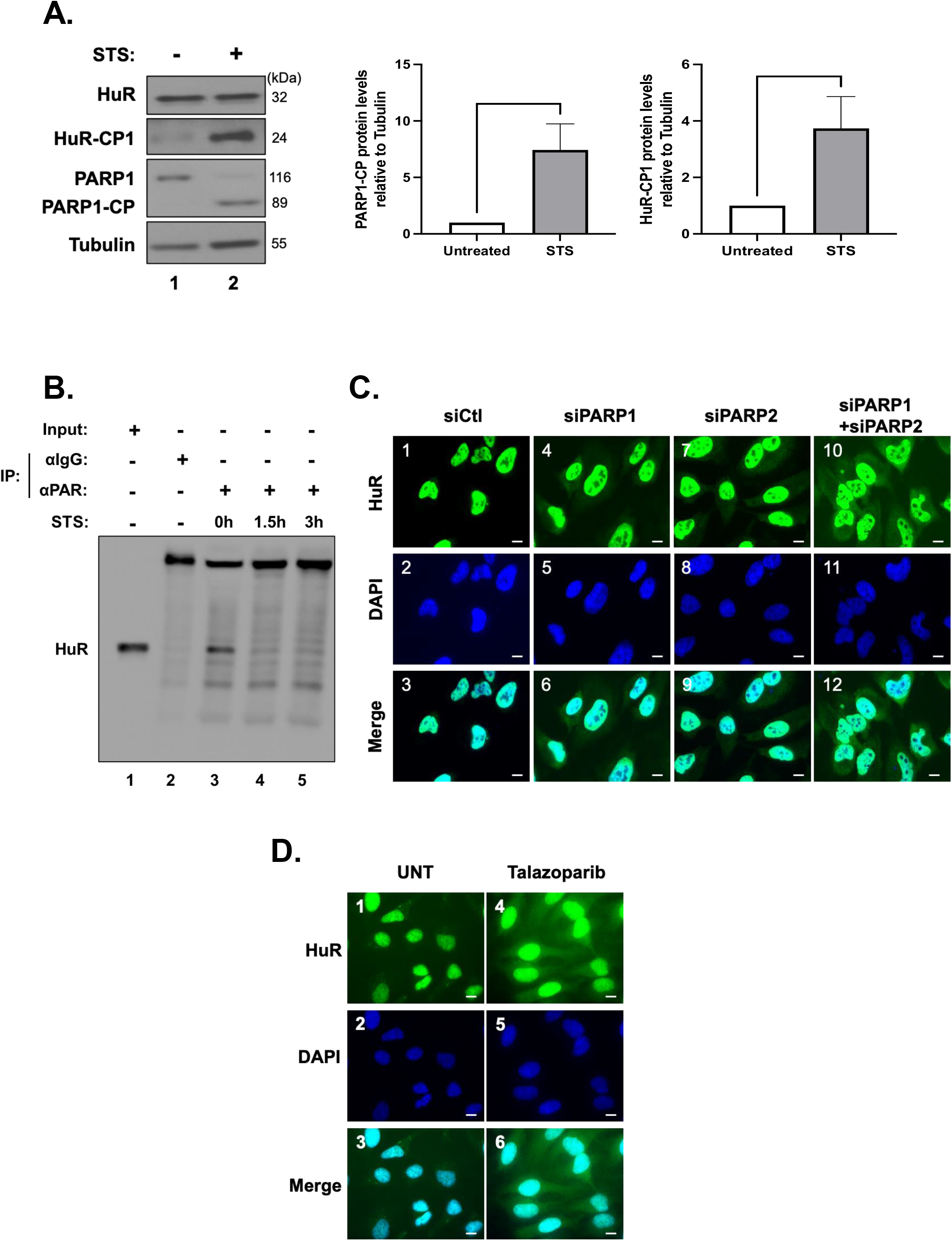
The cytoplasmic translocation and cleavage of HuR in response to an apoptotic stimulus correlates with the cleavage of PARP1/2. **A) (Left)** Hela cells treated with or without 1 µM STS for 1.5 h were collected, lysed, and used for Western blot analysis with antibodies against HuR, PARP1 or ⍺-tubulin (loading control). **(Right)** Densitometric quantification of HuR-CP1 and PARP1-CP signals in the western blot relative to ⍺-tubulin signal. Values were quantified using ImageJ. **B)** Lysates obtained from Hela cells treated with 1 µM STS for (0, 1.5, 3 h) were used for immunoprecipitation experiments using antibodies against PAR or IgG as a negative control. The binding of PAR to HuR was then assessed by western blot using an anti-HuR antibody (3A2). All blots shown in the figure are representative of 3 independent experiments. **C)** HeLa cells were transfected with siRNA targeting PARP1 and/or PARP2 or a control siRNA. These cells were then fixed, permeabilized, and stained with antibodies against HuR. DAPI was used to stain nuclei. Images are representative of three independent experiments. (Scale bars, 10 µm). **D)** Immunofluorescence experiments demonstrating HeLa cells treated with and without 1µm-Talaxzoparib. After 24 hours of treatment, cells were fixed, stained, permeabilized and stained with antibodies against HuR. DAPI was used to stain nuclei. Images of a single representative field are shown and are representation of three independent experiments. (Scale bars, 10 µm). Data presented in Figure 1 are +/- the S.E.M. of three independent experiments with *P<0.05, by unpaired *t*-test.

As mentioned earlier, our lab has shown that in response to lethal stress the accumulation of HuR in the cytoplasm is required for its cleavage and pro-apoptotic function (Supp. Fig.1A) (Mazroui *et al*., 2008). Since the interaction of HuR with PAR only occurs in untreated cells where HuR is localized in the nucleus, we decided to assess if PARP1-mediated PARylation prevents the accumulation of HuR to the cytoplasm. PARP2 has highly redundant function to PARP1, and both are cleaved in a caspase-dependent manner during apoptosis (Ali *et al*, 2016; Benchoua *et al*, 2002). Thus, we sought to assess if depleting both PARPs individually, and in combination, using siRNAs specifically targeting each PARP would affect the localization of HuR (Fig.1C). We observed that these siRNAs efficiently depleted the expression of both PARPs by more than 90% in these cells (Supp. Fig.1B). By performing immunofluorescence experiments, we further observed that although knocking down PARP1 or PARP2 increased the cytoplasmic localization of HuR in untreated conditions, this effect was more prominent when cells were simultaneously treated with siRNAs targeting both proteins (Fig.1C). Additionally, this observation was reproduced using Talazoparib, a well-known PARP1/2 inhibitor (Fig.1D). Our results, therefore, suggest that the interaction of HuR with PAR in the nucleus regulates its nucleocytoplasmic shuttling function under normal conditions in Hela cells.

Next, to determine the impact of PARP1 and PARP2 on the apoptotic function of HuR, we assessed if knocking down these PARPs affects its cleavage. We noticed that HuR cleavage is significantly increased in PARP1 depleted cells under normal conditions (Fig.2A lane 2). We observed, similarly, a trend in HuR cleavage in cells depleted of PARP2 (Fig.2A lane 3). This cleavage, however, was further significantly increased with the double knockdown of PARP1 and PARP2 compared to siCtl treated conditions (Fig.2A lane 4). These results indicate that the PARylation of HuR could play a potential role in modulating its pro-apoptotic function. Since depleting these PARPs resulted in the cleavage of HuR in untreated conditions, mimicking what we observed in the apoptotic conditions, we next questioned the impact of depleting these PARPs on caspase-3 cleavage, another well-established event in apoptosis. As expected, silencing PARP1 and PARP2 together showed a significant increase in the cleavage of caspase-3 (Fig.2A, lane 4). This result was further confirmed by performing flow cytometry experiments which demonstrated a significant increase in the number of Annexin V-positive cells in siPARP1 and siPARP2 treated cells (Fig.2B). Together, these findings highlighted the importance of PARP1/2-mediated PARylation on HuR’s pro-apoptotic function.

**Figure 2:**
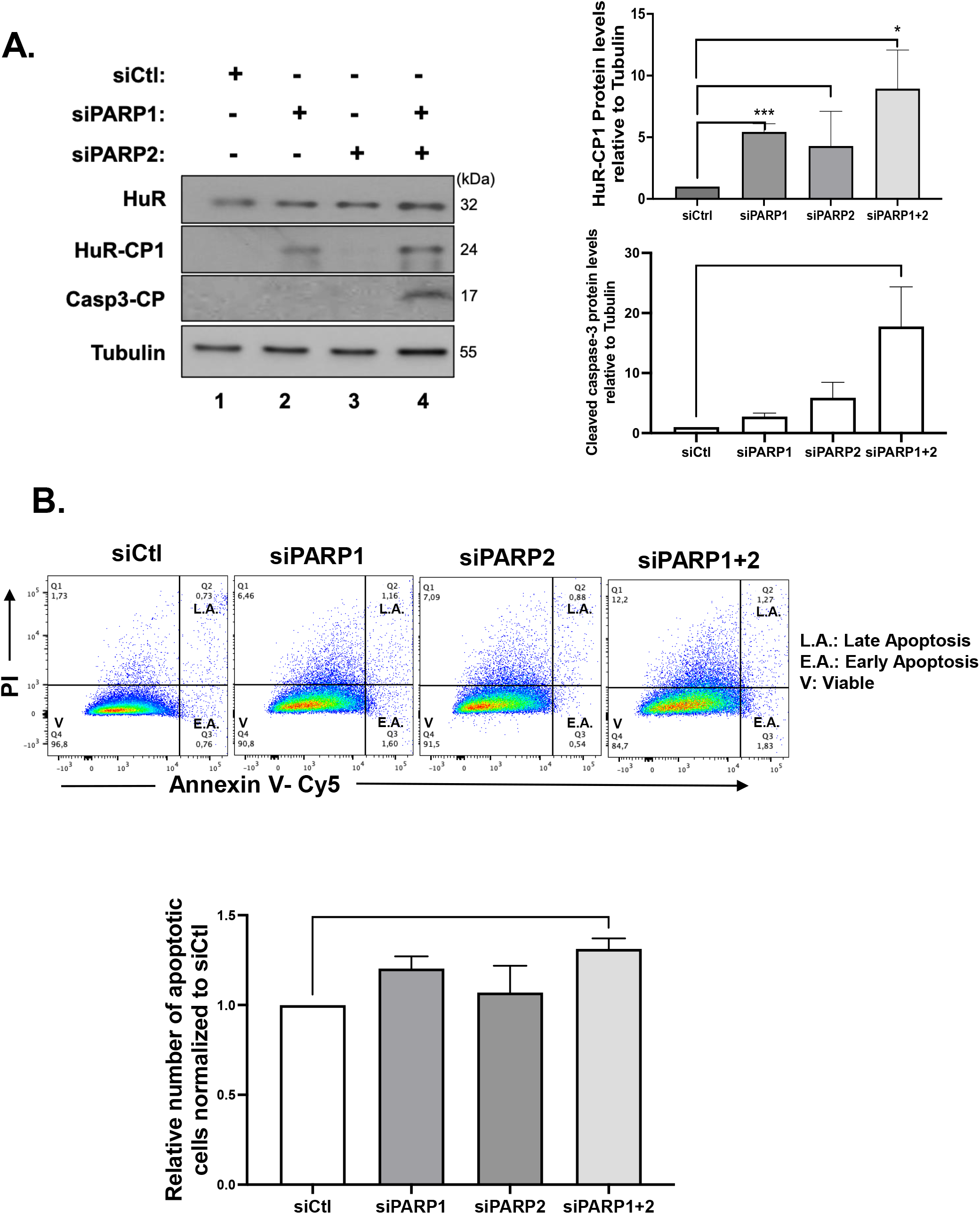
PARP1/2 knockdown increases HuR cleavage and triggers apoptosis. **A)** HeLa cells were transfected with siRNA targeting PARP1 and/or PARP2 or a non-specific control siRNA (siCtl). Lysates were used for Western blot analysis (left panel) with antibodies against HuR, Caspase3 cleavage product (CP) and ⍺-tubulin. Densitometric quantification (Right panel) of HuR-CP1 and Caspase3-CP levels. Values were quantified using ImageJ, normalized to tubulin and shown relative to siCtl. **B)** HeLa cells treated with siRNA as described in A were analyzed by staining with annexin V–Cy5 and PI (Propidium Iodide) and by flow cytometry. The relative number of apoptotic cells was determined for siPARP1 and/or siPARP2. The values are relative to control siRNA treated cells. Data presented in Figure 2 are +/- the S.E.M. of three independent experiments with *P<0.05, ***P<0.001 by unpaired *t*-test.

### PAR binds HuR non-covalently through a consensus motif

It is well-established that PARP-mediated PARylation of target proteins can occur via either covalent modifications or by non-covalent association to PAR(Gagné *et al*, 2008). Both mechanisms were shown to entail different functional consequences on the affected proteins(Duan *et al*, 2019; Ji & Tulin, 2013). Thus, as a first step, we decided to investigate if the non-covalent association of PAR to HuR could mediate its pro-apoptotic function. By performing an *in-vitro* dot blot assay we demonstrated that HuR, unlike BSA and GST (used as negative controls), non-covalently binds to PAR (Fig.3A). Next, we wanted to determine the exact PAR binding site on HuR. To this end, we performed an *in-vitro* peptide mapping experiment where we generated small peptide fragments spanning the complete HuR sequence. Each fragment is about 20 amino acids in length. We found that several fragments (B6, B7, E1) of HuR exhibited binding to PAR with various strength (Fig.3B). However, only one of these (E1) harbours a region (amino acids 201-208 of HuR) that exhibits 76% similarity to a well-known consensus PAR binding site ([HKR]_1_-X_2_-X_3_-[AIQVY]_4_-[KR]_5_-[KR]_6_-[AILV]_7_-[FILPV]_8)_ ^(Gagné^ *^et^ ^al.^*, ^2008; Pleschke^ *^et^ ^al.^*, ^2000)^. Therefore, we dubbed this element as the HuR PAR Binding Site (HuR-PBS).

**Figure 3:**
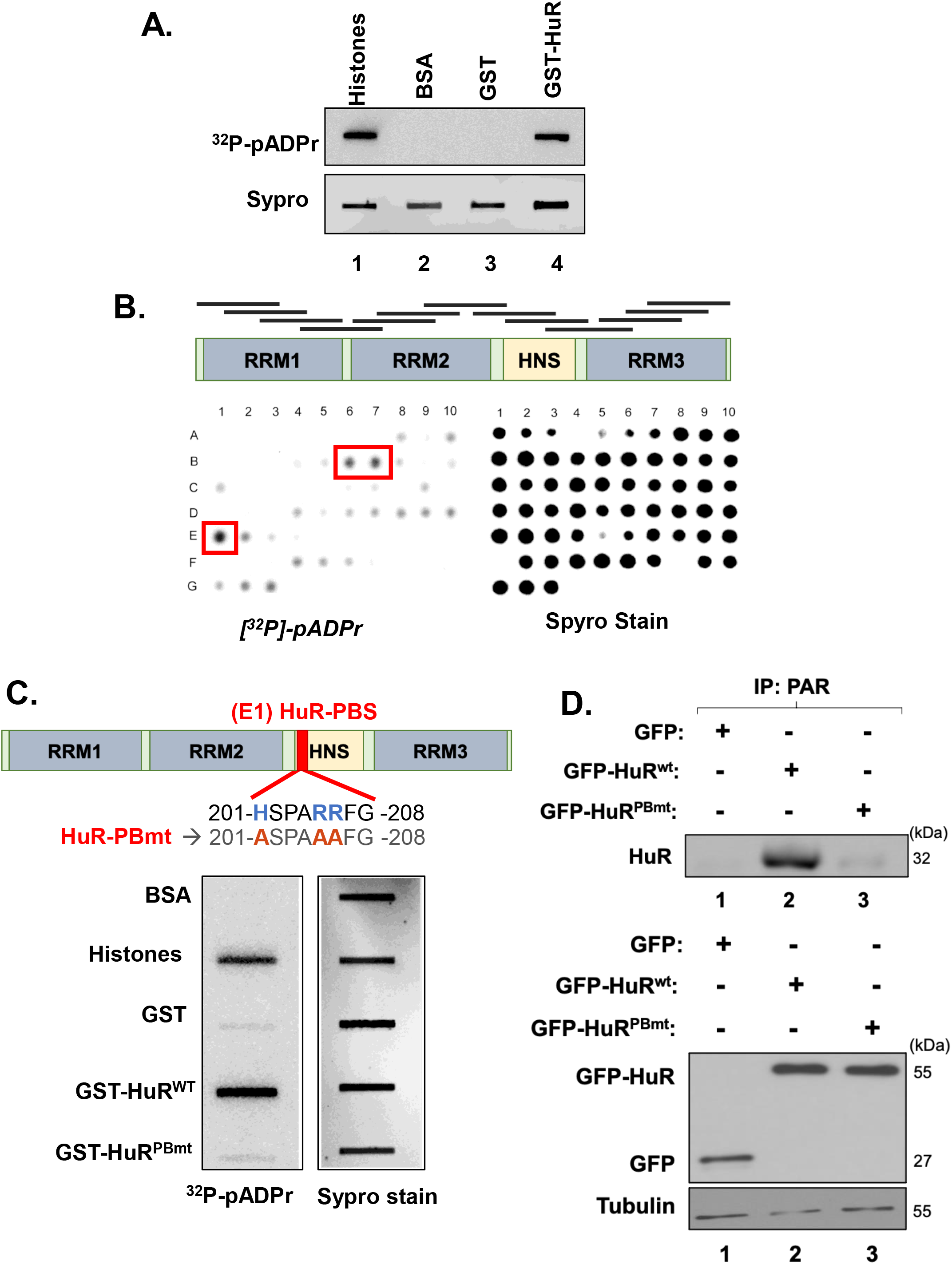
PAR binds HuR non-covalently through a consensus motif. **A)** Recombinant GST and GST-HuR proteins as well as recombinant Histone (positive control) and BSA (negative control) were blotted directly onto nitrocellulose membrane, rinsed, incubated with a radiolabeled ^23^P-pADPr and analyzed by autoradiography. Sypro Ruby stain was used to demonstrate the integrity and quantity of the proteins. **B)** HuR protein was fragmented into 63 small peptides (each fragment is 20 amino acids in length with 5 staggered amino acids) used for peptide mapping experiment. All fragments were blotted onto nitrocellulose membrane and processed as in A. **C)** (**Top**) Schematic showing location of PAR binding site of HuR. Mutation of this site was generated by substituting the positively charged amino acid Histidine and Arginine by hydrophobic Alanines residues. (**Bottom**) Slot blot assay was performed using recombinant GST-HuR^wt^, GST-HuR^PBmt^ protein and GST/BSA as a negative control, while Histone as a positive control. Sypro Ruby stain was used to demonstrate the integrity and quantity of the proteins. **D)** (**Top**) Total cell extracts obtained from HeLa cells transfected with GFP, GFP-HuR^wt^ or GFP-HuR^PBmt^ were used for immunoprecipitation experiments using antibodies against PAR. The binding of PAR to HuR was then assessed by western blot using anti-HuR antibody. (**Bottom**) Transfection efficiency was assessed by determining the levels of these proteins in the input using anti-GFP and ⍺-tubulin as a loading control. Immunoprecipitation results are representative of three independent experiments.

To better understand the importance of this site on the function of HuR, we generated a mutant isoform of HuR (HuR^pbmt^) whereby the (+) charged arginines (R) and histidines (H) residues within the HuR-PBS were converted into alanines (A) (Fig.3C top). Using the dot-blot approach mentioned above we demonstrated that unlike the wild type HuR (HuR^wt^), the HuR PAR binding mutant (HuR^pbmt^) lost its ability to bind PAR (Fig.3C bottom). We next assessed the importance of this site for PAR binding in HeLa cells by transfecting the cells with GFP, GFP-HuR^wt^ and GFP-HuR^pbmt^ followed by a co-IP experiment where we immunoprecipitated PAR and immunoblotted with anti-HuR (Fig.3D). We demonstrated that although PAR binds to GFP-HuR^wt^ this interaction is completely abrogated due to mutation of the PAR-binding site (Fig.3D). Together, these results reveal that HuR non-covalently interacts with PAR through the harbored PAR-binding motif.

### PAR binding prevents the pro-apoptotic function of HuR by promoting its nuclear localization

Our results described above show that the depletion of PARP1/2 decreased the nuclear localization of HuR. This is likely due to the decreased interaction of HuR to PAR. To assess if this is the case, we next assessed the impact of mutating the HuR PAR-binding site on its cellular localization. Immunofluorescence assays revealed that HuR^Pbmt^ but not HuR^wt^ accumulates in the cytoplasm of untreated HeLa cells, mimicking the observations obtained due to the knockdown of PARP1/2 (Fig.4A). Together our results therefore suggest that the non-covalent association of PAR with HuR plays an important role in modulating its cellular localization in normal HeLa cells.

**Figure 4:**
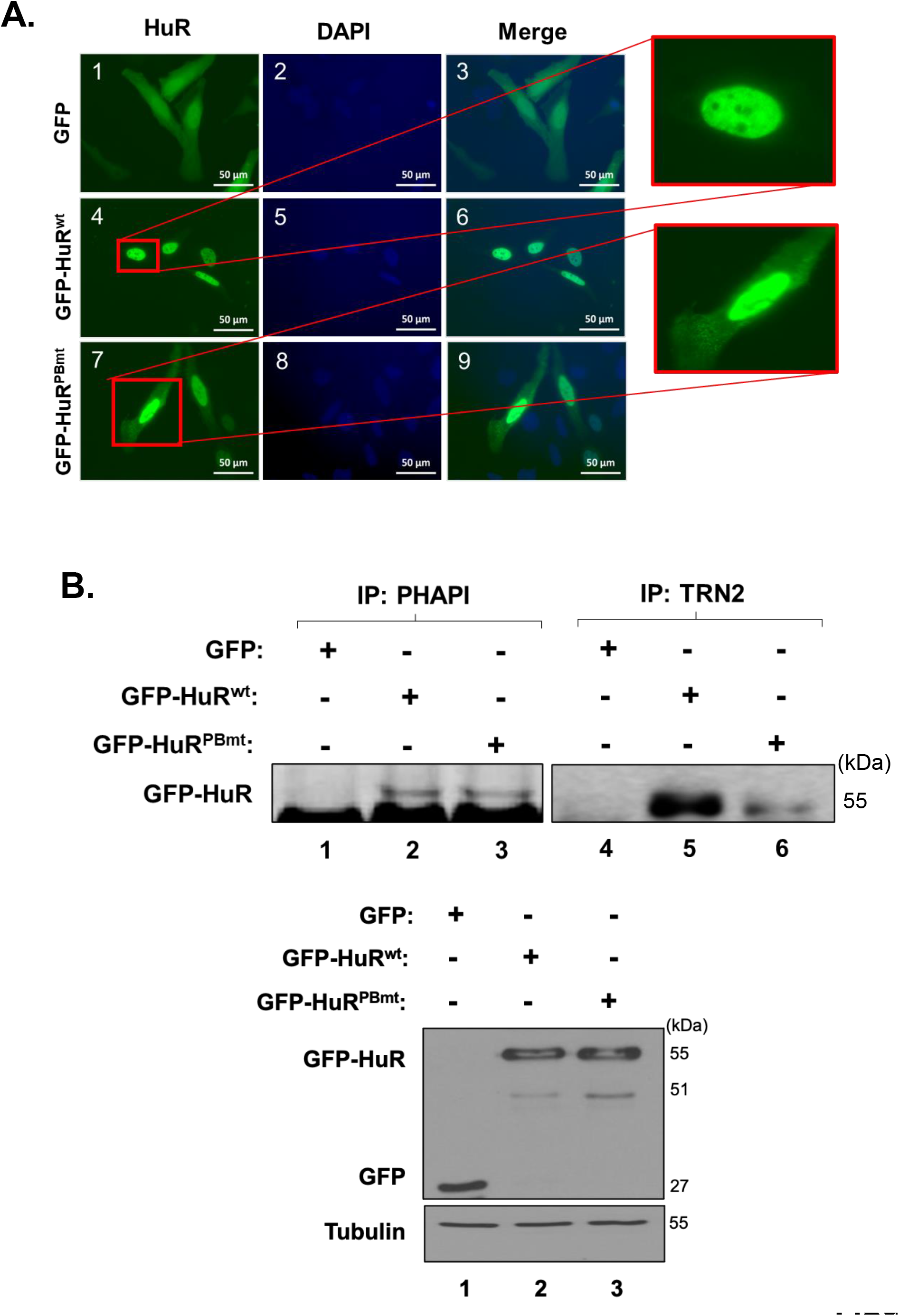
HuR binding to PAR modulates its cellular localization in Hela cells. **A)** HeLa cells transfected with GFP, GFP-HuR^wt^ and GFP-HuR^PBmt^ were fixed, permeabilized, and stained with antibodies against HuR and DAPI. Images are representative of 3 independent experiments. (Scale bars, 50 µm). **B)** Total cell extracts obtained from HeLa cells transfected as in A were used for immunoprecipitation experiments using antibodies against PP32/PHAPI (left panel) or TRN2 (right panel). Immunoprecipitated complex was then assessed by western blot using HuR antibodies. The blots are representative of 3 independent experiments.

We and others have shown that the nucleocytoplasmic translocation of HuR, during apoptosis, is mediated by its association with adaptor proteins for nuclear export such as PHAP-I and with the import factors such as transportin-2, (Brennan *et al*, 2000; Mazroui *et al*., 2008; Zhang *et al*, 2016). To determine whether mutating the PAR binding site would have an impact on the differential association of HuR with these proteins, we immunoprecipitated PHAPI and TRN2 individually and assessed their association with GFP-HuR^wt^ or GFP-HuR^Pbmt^ (Fig.4B). We observed that, unlike HuR^wt^, the HuR^Pbmt^ isoform loses its association with TRN2 (Fig.4B right) but not with PHAPI (Fig.4B left). This finding suggests that an intact HuR-PBS is required for the association of HuR with TRN2 and its retention in the nucleus. We have previously shown that the cleavage of HuR is tightly related to its cytoplasmic accumulation, due to the competition of HuR-CP1 with full length HuR for the binding to TRN2, leading to the accumulation of full length HuR in the cytoplasm. Therefore, we next determined whether mutating the HuR-PBS would affect the cleavage of HuR. We observed that the GFP-HuR^Pbmt^ is cleaved to a greater extent than GFP-HuR^wt^(Fig.5A). Interestingly, we observed that the expression of HuR^Pbmt^ increased the cleavage of caspase-3 to a greater extent than cells expressing HuR^wt^ (Fig.5A). To determine the physiological importance of PAR binding, we performed flow cytometry analysis to assess the cell fate of HuR^Pbmt^ expressing cells compared to cells expressing HuR^wt^. These results further supported our findings described above and showed an increase in annexin V positive cells expressing HuR^pbmt^ (Fig.5B), providing evidence for the anti-apoptotic role conferred to HuR by the binding to PAR. In conclusion, we demonstrate that HuR can non-covalently bind to PAR, and that this binding alters its pro-apoptotic function through the regulation of its localization, further underlining the importance of HuR localization in its pro-apoptotic function.

**Figure 5:**
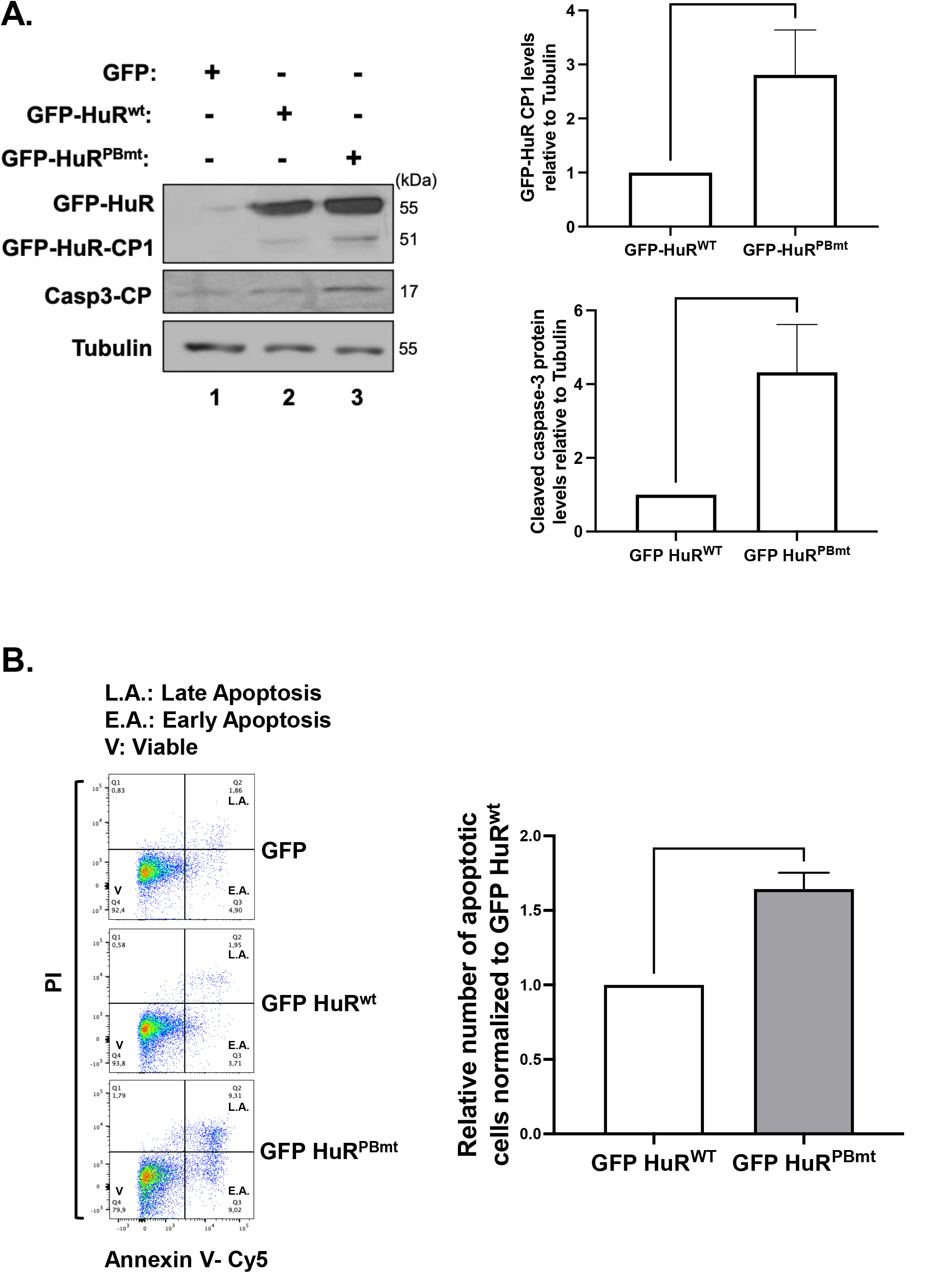
PAR binding to HuR negatively affect its pro-apoptotic function. **A)** HeLa cells were transfected withGFP, GFP-HuR^wt^ and GFP-HuR^PBmt^. (**Left**) Lysates were used for Western blot analysis with antibodies against HuR, Caspase3-CP and ⍺-tubulin. (**Right**) Densitometric quantification of HuR-CP1 and Caspase3-CP levels were normalized to ⍺-tubulin level and shown relative to GFP-HuR^wt^. **B)** HeLa cells transfected as described in A were analyzed by staining with annexin V–Cy5 and PI and analyzed by flow cytometry. The relative number of apoptotic cells was determined for GFP-HuR^wt^ and GFP-HuR^PBmt^ transfected HeLa cells. The values were normalized to GFP. Data presented in Figure 5 are +/- the S.E.M. of three independent experiments with *P<0.05, ***P<0.001 by unpaired *t*-test.

## Discussion

In this study we identify PARylation as a regulatory mechanism that modulates the function of HuR in determining cell fate. Our results show that PARP1/2-mediated PARylation prevents the accumulation of HuR to the cytoplasm resulting in a decrease in its cleavage, and inhibition of HuR’s pro-apoptotic function. We demonstrated that the combined depletion of PARP1 and PARP2 increases the cytoplasmic accumulation of HuR and thus increases its cleavage. HuR cleavage, consequently, increases its pro-apoptotic function as evidenced by the significant increase in the level of caspase-3 cleavage and in the number of apoptotic cells. Furthermore, we showed that PAR binds HuR non-covalently through a consensus motif and that this binding is required for the nuclear localization of HuR as well as its association with the import factor TRN2. Indeed, we found that mutating HuR-PBS prevented PAR from binding to HuR resulting in the cytoplasmic accumulation of HuR and therefore advancing apoptosis. Thus, our work provides evidence for the importance of the PARP-mediated PARylation and the resulting PAR binding to HuR in regulating the function of HuR during apoptosis (Fig. 6).

**Figure 6:**
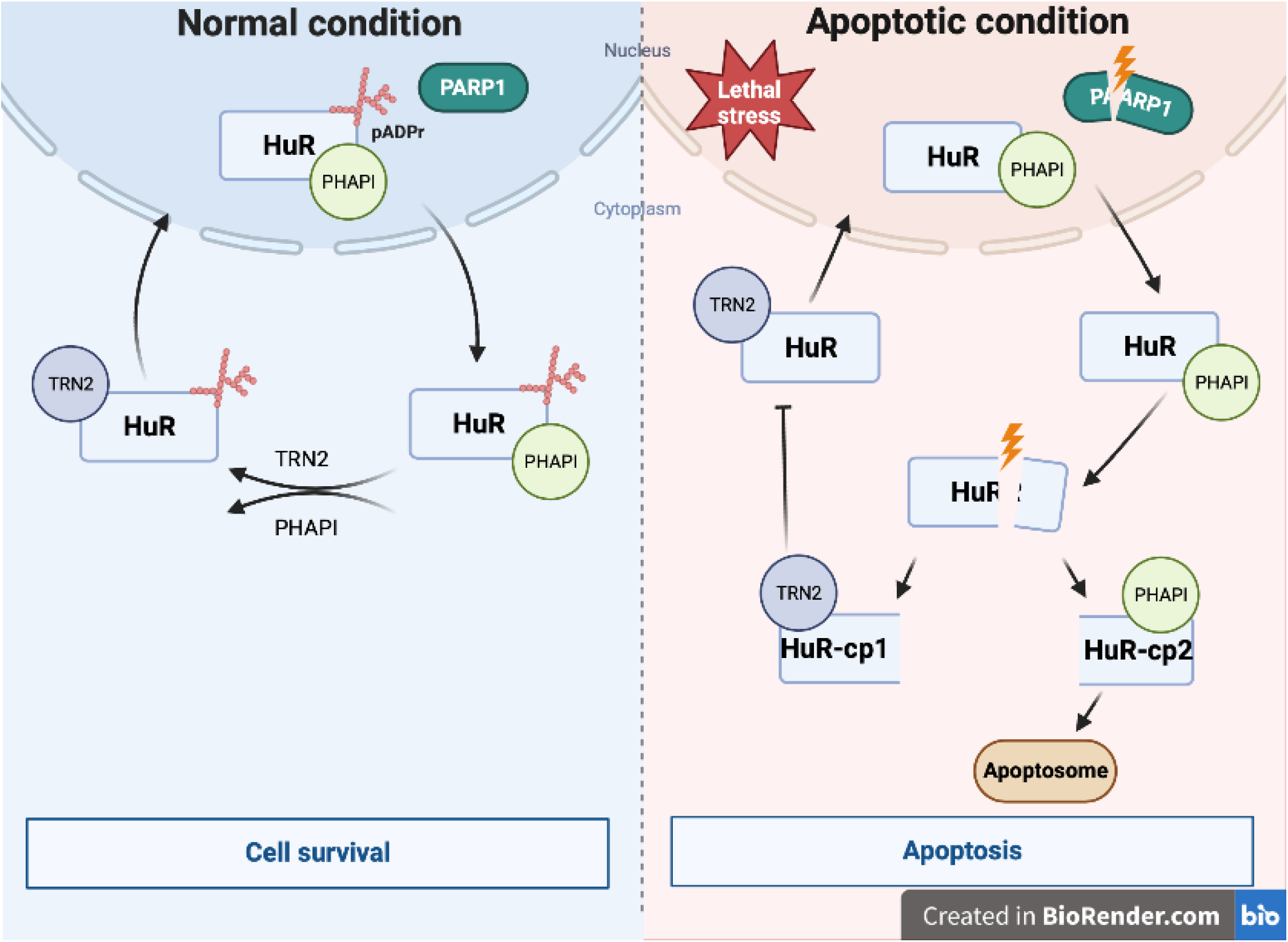
Proposed model. Model depicting the mechanism by which HuR association with PAR polymers regulates its apoptotic function. Under normal condition, HuR interacts with PAR polymers through its PAR Binding Site (HuR-PBS) maintaining its nuclear localization by promoting its interaction with the import factor TRN2. In response to a lethal assault, HuR loses its binding to PAR concurrently with the cleavage of PARP1. HuR/PHAPI translocate to the cytoplasm where HuR undergoes caspase mediated cleavage yielding HuR-CP1 and HuR-CP2. While HuR-CP2 interact with PHAPI mediating the activation of apoptosome-formation, HuR-CP1 interacts with TRN2 preventing the reuptake of HuR back to the nucleus. HuR, therefore, accumulates in the cytoplasm, advancing apoptosis.

Although PARylation of HuR has been previously shown to regulate the function of HuR during inflammation(Ke *et al*., 2017; Ke *et al*, 2021) as well as muscle cell differentiation (Mubaid *et al*. unpublished data), the importance of this modification on HuR function in cell fate was not assessed. Indeed, recent studies by Ke *et. al* revealed that, in response to inflammatory stimuli, PARP1-mediated PARylation of HuR binds to its HNS-RRM3 region and modifies it at a conserved aspartate residue D226. They demonstrated that mutating this site (D226) or inhibiting PARP impacted HuR localization, its ability to associate with pro-inflammatory messages as well its oligomerization (Ke *et al*., 2017; Ke *et al*., 2021). More recently, our lab uncovered that PARylation of HuR by Tankyrase1 (TNKS1), also known as PARP5a, promoted HuR cytoplasmic accumulation and cleavage as well as its ability to associate with promyogenic mRNA during myogenesis (Mubaid *et al*. unpublished data). In this study, however, we identified a consensus PAR binding motif located within the HNS of HuR, and we showed that PARP1/2-mediated PARylation and PAR binding to HuR through this identified motif mediates its subcellular localization and function during apoptosis.

Although HuR is predominantly a nuclear protein under normal conditions, its HNS domain encompasses a nucleocytoplasmic shuttling sequence allowing it to shuttle between the nucleus and the cytoplasm in response to various stimuli, such as stress signals (Fan & Steitz, 1998; Grammatikakis *et al*., 2017a; van der Giessen & Gallouzi, 2007; Von Roretz *et al*., 2013). This translocation is important for the HuR-mediated post-transcriptional regulation of many mRNA targets including mRNA localization, stabilization, and translation, and has been shown to have physiological relevance by affecting cell fate and muscle cell differentiation (Brennan & Steitz, 2001; Mazroui *et al*., 2008; van der Giessen & Gallouzi, 2007; von Roretz *et al*, 2011a). Several studies have reported that post-translational modification (PTMs) of residues within the RRMs influenced the function of HuR in regulating RNA metabolism, while modification of residues within or near the HNS impacted HuR subcellular localization (Gallouzi & Steitz, 2001; Grammatikakis *et al*., 2017a; Srikantan & Gorospe, 2012). For example, phosphorylation of HuR by Chk2 at HuR residues S88, S100 and T118 located within RRM1 and RRM2 modulates HuR binding to SIRT1 mRNA and other mRNA targets (Abdelmohsen *et al*, 2007). On the other hand, phosphorylation by cdk1 at S202 facilitated HuR binding to the nuclear 14-3-3 triggering its nuclear retention (Grammatikakis *et al*., 2017a; Kim *et al*, 2008). Previously, our lab and others have shown that the localization of HuR is dependant of its HNS, which mediates the differential association of HuR with protein partners for nuclear export, such as PHAP-I and APRIL, and with the import factors TRN-1-2, and importin α (Brennan *et al*., 2000; Mazroui *et al*., 2008; Rebane *et al*, 2004). It has been shown that, under normal condition, HuR is localized mainly to the nucleus (Mazroui *et al*., 2008). However, in response to lethal stress, HuR and PHAPI translocate to the cytoplasm where it is cleaved in a PKR-dependent pathway by caspase-3 and -7 yielding two cleavage products (HuR-CP1 and HuR-CP2) (Beauchamp *et al*, 2010; Mazroui *et al*., 2008; von Roretz & Gallouzi, 2010b; Von Roretz *et al*., 2013). Moreover, our lab showed that HuR-CP1 associate with TRN2 preventing the nuclear reuptake of HuR, thus causing HuR to accumulate in the cytoplasm(von Roretz *et al*., 2011a). In this present study, we identified the non-covalent binding of PAR to HuR as a regulatory mechanism mediating its association with these partners and therefore, its pro-apoptotic function. Given that the HuR-PBS is located in the HNS, it is not surprising that it regulates the localization of HuR. Our results demonstrate that an intact PAR binding to HuR is required for its binding with TRN2 in particular, and that mutating this site resulted in the loss of this binding. As HuR binding to different protein partners seems to influence the function of HuR during apoptosis, it would be valuable to determine if mutating the PBS on HuR would have an impact on HuR association with different protein ligands in other cell systems. Also, as previous observations highlighted the importance of HuR cleavage products during the onset of apoptosis, it would be interesting to investigate the role of PARP mediated modification on HuR and/or binding to PAR on the role of HuR-CPs as pro-apoptotic players.

In addition to the ability of PARylation to covalently modify acceptor protein at specific residues, a number of proteins, also known as PAR readers, can be modified by the non-covalent attachment of PAR polymers to consensus PAR binding motifs [16, 28]. In fact, many RBPs have been shown to be bound by PAR covalently and non-covalently, both of which lead to the alteration of their functions [16]. Previous studies have reported that HuR is covalently PARylated by PARP1 at the D226 residue in LPS-induced cells, which affected its localization and function(Ke *et al*., 2017; Ke *et al*., 2021). Moreover, HuR also has been shown, in a proteome-wide analysis of PAR-associated proteins, to bind PAR non-covalently(Jungmichel *et al*, 2013). In our study we confirmed that PAR binds HuR non-covalently and we identified the consensus PAR binding site and showed its physiological importance in the anti-apoptotic function of HuR. Although the covalent PARylation of HuR at the D226 residue is critical in regulating its function and localization in macrophages(Ke *et al*., 2017), our results indicate that this modification is likely not involved in regulating HuR function in normal HeLa cells. Indeed, we observed that mutating the PAR-binding site on its own was sufficient to completely inhibit the pulldown of HuR due to the immunoprecipitation of PAR from normal Hela cells (Fig. 3D) and that, furthermore, the HuR-PAR binding mutant is localized to the cytoplasm in these cells where it is cleaved (Figs. 4, 5).

Many recent studies are now pointing to the importance of these two manners of PARylation on the function of their substrates and how there may be an interplay between the two modifications(Alemasova & Lavrik, 2019; Grammatikakis *et al*., 2017a; Ke *et al*, 2019). For instance, heterogeneous nuclear ribonucleoprotein A1(hnRNPA1), a well-known RBP, has been shown to be PARylated covalently and it can also bind to PAR or PARylated proteins non-covalently. Recently, Duan *et al*. showed that hnRNPA1 is PARylated on Lysine 298 and mutating this site decreased its PARylation level and affected its localization(Duan *et al*., 2019). They also showed that when the PAR binding motif is mutated, it increased its covalent PARylation (Duan *et al*., 2019). These observations led them to suggest that the non-covalent PAR binding reduces the hyper-PARylation of hnRNPA1. Therefore, it would not be surprising that such an interplay exists between the covalent and non-covalent PARylation of HuR during apoptosis and potentially other systems. Importantly, this study shows that mutating the non-covalent PARylation binding site prevented oligomerization of hnRNPA1 and prevented the formation of stress granule(Duan *et al*., 2019). This impact is not surprising, since PARylation is suggested to nucleate membranelles organelles, including stress granules (Alemasova & Lavrik, 2019; Isabelle *et al*, 2012; Ke *et al*., 2019). HuR has been shown to locate in membranelles organelles and is well known to form oligomers, which might be potentially regulated by PARylation, similar to hnRNPA1(Isabelle *et al*., 2012). It is thus possible that the non-covalent binding of HuR can prevent, as was shown for hnRNP A1, its covalent modification by PARPs under normal conditions. In doing so, this event may explain the differential role of HuR in modulating the survival or death of cells under normal or stress-induced conditions.

Our work, thus, has furthered our understanding of the role of HuR in apoptosis, showing that it is regulated by PARylation. Moreover, understanding the regulatory mechanism underling the pivotal role of HuR in cell fate will bring a new hope to find therapies to overcome many diseases, such as numerous cancers, that are associated with the increased cytoplasmic localization of HuR.

## MATERIAL AND METHODS

### Cell culture, transfection, and treatment

HeLa CCL-2 cells (American Type Culture Collection) were grown and maintained in DME (Dulbecco’s modified Eagle) media (Invitrogen) containing 10% FBS (Sigma) and 1% penicillin/streptomycin (Sigma). Plasmid and siRNA were transfected as described by the Polyplus jetPRIME transfection protocol using 0.5ug/mL and 50nM/mL plasmid and siRNA, respectively. Plasmid transfection was done on 80% confluent HeLa cells, whereas siRNA transfection was done on 60% confluent HeLa cells. siRNAs were purchased from Ambion: siPARP1 (ID: s1097), siPAPR2 (ID: 111561). For the STS treatment (Sigma Aldrich), HeLa cells were incubated in 1uM STS for 1.5 or 3 hours. Treatments were done 24 hours post transfection. For PARP inhibitors experiment, cells were treated with Talazoparib 1 uM from Selleckchem (BMN673) for 24 hours.

### Plasmid construction and protein purification

The GFP-HuR^wt^ and GST-HuR plasmids were generated as described (Mazroui *et al*., 2008). The GFP-HuR^pbmt^ and GST-HuR^pbmt^ plasmids were generated (by mutating the Histidine and Arginine amino acids to Alanines) by Norclone Biotech Laboratories (London, ON, Canada). The GST, GST-HuR^wt^ and GST-HuR^pbmt^ recombinant proteins were generated by transforming BL21 with the respective plasmids. The expression of the proteins was induced by IPTG (0.5mM for 4hours at 37°C) in a 1-liter culture. The bacteria were collected and lysed. The GST proteins were pulled down using Glutathione Sepharose beads and processed as previously described (Brennan *et al*., 2000).

### Protein extraction and immunoblotting

Total cell extract from HeLa cells were prepared as described previously (von Roretz & Gallouzi, 2010a). Briefly, cell extracts were lysed with mammalian lysis buffer (50 mM HEPES pH 7.0, 150 mM NaCl, 10% glycerol, 1% Triton, 10 mM pyrophosphate sodium, 100 mM NaF, 1 mM EGTA, 1.5mM MgCl_2_, 1 X protease inhibitor (Roche) and 0.1 M orthovanadate), then lysates were collected after centrifugation for 15 minutes at 12000rpm. Western blotting was performed as described (Mazroui *et al*., 2008) using the following antibodies: HuR (3A2 (Gallouzi *et al*, 2000), 1:1000), α-tubulin (Developmental studies Hybridoma Bank, 1:10000), Cleaved Caspase-3 (Cell signaling, 1:1000), full length PARP (Cell signaling, 1:1000), GFP (JL-8, Living colors, 1:1000). Quantification of bands on Western was done using ImageJ (Fiji) software and normalized to α-tubulin. Statistical analysis for significance was performed using GraphPad software with a one-tailed unpaired *t-*test.

### Binding (Dot/Slot blot) assay and peptides mapping experiments

These experiments were performed as described (Pleschke *et al*., 2000). Briefly GST, GST-HuR^wt^, GST-HuR^pbmt^, Histone (positive control) and BSA (negative control) or peptides spanning the HuR protein (fragmented into 63 peptides; each is 20 amino acids in length) were dot-blotted directly onto nitrocellulose membrane. The blot was then rinsed three times with TBST (Tris-buffered saline with 0.1% Tween 20 detergent) and incubated with radioactive pADPr (^32^P-pADPr) generated by auto-modified PARP1, washed, and probed for retention of the pADPr. After incubation for 1 hour at room temperature with gentle agitation, the membrane was washed, dried, and subjected to autoradiography. Peptides/ full length proteins were incubated with Sypro Ruby stain in order to demonstrate their integrity and event distribution.

### Immunoprecipitation

Immunoprecipitation experiments were performed as previously described (Cammas *et al*, 2014; van der Giessen & Gallouzi, 2007). Briefly, antibodies against anti-PAR 10H clone (Tulips), anti-TRN2 (van der Giessen & Gallouzi, 2007), and anti-PP32/PHAPI (Santa Cruz) were incubated with 60 µl of protein A-Sepharose slurry beads (GE Healthcare) (washed and equilibrated in cell lysis buffer) for 4h at 4 °C. Beads were washed three times with cell lysis buffer (25mM Tris-HCl pH 8.0, 650mM NaCl, 0.05% Tween-20, 100mM NaF, 1X protease inhibitors) and then incubated with 800 µg of total cell extracts (TCE) overnight at 4 °C. Beads were subsequently washed three times with cell lysis buffer and Lamelli dye was added to the immunoprecipitated proteins for analysis by western blot.

### Immunofluorescence

Immunofluorescence was performed as previously described (von Roretz *et al*., 2011a). Cells were fixed in 3% Paraformaldehyde (Sigma) for 20min and then permeabilized with 0.1% Triton-X 100 in PBS-goat serum for 20 min. After permeabilization, cells were incubated with primary antibodies against HuR/3A2 (1:1500) in 1% normal goat serum/PBS at room temperature for 1 hr. The cells were then incubated with the secondary antibody (Alexa Fluor® 488) and 4’,6-diamidino-2-phenylindole (DAPI) (for nuclear staining). Images were taken at room temperature with a 63X oil objective on an inverted Axiovert 200M microscope with an Axiocam MRm digital camera (Zeiss).

### Annexin V–Cy5/PI assay

Cells in the tissue culture dishes was collected 24h post-transfection by trypsinizing the plate. Cell pellets were then processed as described by the apoptosis detection reagent kit protocol (Abcam; ab14147). Apoptotic and necrotic cells were identified by annexin V–Cy5 and Propidium Iodide (PI) staining, respectively, using the flow cytometry analyzer (FACSCanto II). The flow cytometry work was performed in the Flow Cytometry Core Facility for flow cytometry and single cell analysis of the Life Science Complex. The data analysis was then analyzed using FlowJo software.

### Quantitative RT-qPCR

RNA was extracted from cell extracts using Trizol reagent (Invitrogen) according to the manufacturer’s instructions. One microgram of total RNA was reverse transcribed using the 5X iScript reagent (Bio-Rad) according to the manufacturer’s protocol. Each cDNA sample was diluted 20 folds and used to detect the mRNA levels of *PARP1, PARP2* and *GAPDH (*used as a loading control) using the SsoFast EvaGreen reagent (Bio-Rad Laboratories). The relative expression level was calculated using the 2^-ΔΔCt^ method, where ΔΔCt is the difference in Ct values between the target and reference genes (GAPDH). Primers used for qPCR are as follows. PARP1 (F: 5’-CCC AGG GTC TTC GAA TAG-3’, R: 5’-AGC GTG CTT CAG TTC ATA C-3’), PARP2 (F: 5’-GGA AGG CGA GTG CTA AAT GAA-3’,R: 5’-AAG GTC TTC ACA GAG TCT CGA TTG-3’), GAPDH (F: 5’ - AAG GTC ATC CCA GAG CTG AA - 3’, R: 5’ - AGG AGA CAA CCT GGT CCT CA – 3’).

## Data Availability

Any additional information required to reanalyze the data reported in this paper is available from the corresponding contact upon request.

## Acknowledgements

**KA** contributed to conceptualization, conducted the investigation and validation of experimental findings, wrote the original draft, and performed the formal analysis and visualization of experimental findings. **DTH** and **SM** contributed to the conceptualization and conducted the preliminary investigation and validation of experimental findings. **XJL** and **SB** helped by performing some of the Western Blot and qPCR experiments. **SDM and SK** assisted with conceptualization, data analysis, and helped edit and review the manuscript. **JPG** and **GGP** performed the *in vitro* PAR binding assays and peptide mapping experiments and helped with the analysis of the data. **IEG** conceptualized, established, and directed the execution of the project, interpreted the data, reviewed, and edited the manuscript.

---

**Supplemental Figure 1**

**A)** Immunofluorescent experiments demonstrating the localization of HuR in Hela cells treated with or without 1 µM STS for 1.5 h.

**B)** Total RNA was isolated from HeLa cells transfected with siRNA targeting PARP1 and/or PARP2 or a control siRNA and RT-qPCR analysis was performed using primers for PARP1 and PARP2 to determine mRNA level for the validity of knockdown efficiency.

